# FAIM Inhibits Insulin Amyloidogenesis through a Noncanonical Aggregation Pathway

**DOI:** 10.64898/2026.07.13.738277

**Authors:** Dana Wolfe, Jhinuk Saha, Joshua Mitchell, Sam McCalpin, Michael Gutknecht, Charles L. Brooks, Thomas L. Rothstein, Ayyalusamy Ramamoorthy

**Author notes:** Authors for correspondence:] Ayyalusamy Ramamoorthy; Thomas L. Rothstein.

## Abstract

Insulin can misfold and assemble into amyloid fibrils, a process linked not only to complications of insulin therapy but also to proteotoxic stress in pancreatic β-cells. Despite growing interest in the pathological consequences of insulin aggregation, prevention efforts are limited by an incomplete understanding of the endogenous mechanisms that counteract it. Here, we identify Fas apoptosis inhibitory molecule (FAIM) as an endogenous suppressor of insulin amyloid formation. FAIM reduces β-sheet formation and redirects insulin toward disordered, growth-incompetent assemblies. Further, FAIM attenuates the cytotoxicity of insulin aggregates *in vitro*. We hypothesize that this effect arises from masking aggregation-prone regions of insulin and show through structural modeling that FAIM interacts with both insulin chains. These findings extend the anti-aggregation function of FAIM to insulin and suggest a mechanism for endogenous suppression of insulin amyloid formation. More broadly, our results provide insight into the regulation of insulin assembly and highlight FAIM as a candidate modulator of proteostasis in metabolic disease.

**Statement for a broader audience:** Insulin can clump together into harmful aggregates, contributing to complications of insulin therapy and potentially damaging the insulin-producing cells of the pancreas. This study identifies the naturally occurring protein FAIM as a protective factor that inhibits the formation of these harmful aggregates and reduces their toxicity. These findings improve our understanding of how cells protect insulin from harmful aggregation and may open new avenues for developing therapies to combat diabetes-related protein aggregation.

## Introduction

Type 2 diabetes (T2D) is a leading metabolic disease affecting an estimated 590 million individuals worldwide.^1^ The pathogenesis of the disease involves the progressive deterioration of pancreatic β-cell function, resulting in impaired insulin secretion and chronic hyperglycemia.^2–4^ A defining feature of T2D is insulin resistance in peripheral tissues, including skeletal muscle, adipose tissue, and the liver, which increases the demand for insulin secretion.^5^ To compensate, pancreatic β-cells undergo hypersecretion of insulin, increasing demands on protein folding and secretory pathways. These demands place particular importance on maintaining the stability of insulin, the principal hormone responsible for glucose homeostasis.

Insulin is a peptide hormone composed of an A-chain (21 residues) and a B-chain (30 residues) connected by two interchain and one intrachain disulfide bonds.^6,7^ In the secretory granules of β-cells, insulin is stored at high concentrations as zinc-stabilized hexamers and released as bioactive monomers in response to elevated glucose.^8^ However, insulin has an intrinsic propensity to form fibrillar aggregates under conditions including low pH, elevated temperature, and high local concentration. Consistent with this behavior, insulin amyloidosis has been reported in pharmaceutical formulations, at sites of repeated subcutaneous injection, and under experimental conditions that promote protein misfolding.^9–11^ The biological and therapeutic relevance of insulin has led to a substantial body of research into the mechanisms of insulin aggregation.

Through the use of biophysical and cellular approaches, the kinetics, structural transitions, and cytotoxic effects of insulin aggregation have been described in considerable detail.^12–17^ These studies established that insulin readily adopts amyloid structures under destabilizing conditions. As a result, numerous stabilization strategies have been developed to preserve insulin during storage and administration. Formulation approaches commonly employ zinc ions and phenolic preservatives to maintain insulin in its hexameric state, whereas protein engineering has produced analogs such as Lispro and Aspart with reduced tendencies toward self-association and fibrillation.^18,19^ Additional modifications, including PEGylation and Fc-fusion, have also been used to improve solubility and formulation stability.^20^ Despite these advances, efforts to limit insulin aggregation have focused largely on extrinsic stabilization. By contrast, relatively little is known about the endogenous factors that regulate insulin aggregation or the mechanisms by which cells maintain insulin proteostasis.

One candidate regulator of protein aggregation is fas apoptosis inhibitory molecule (FAIM), a small and highly conserved protein originally identified as an inhibitor of fas-mediated apoptosis.^21^ FAIM exists as two major isoforms: the ubiquitously expressed FAIM-S (20 kDa) and the neuron-specific FAIM-L (22 kDa).^22,23^ FAIM lacks significant sequence homology to classical molecular chaperones and does not possess ATPase domains or canonical amyloid-binding motifs.^24,25^ Instead, structural studies by NMR and X-ray crystallography revealed a double clamshell-like β-sandwich with flexible loops extending from the core.^26,27^ This arrangement may facilitate interactions with exposed hydrophobic surfaces on misfolded proteins. FAIM has been shown to suppress aggregation of several amyloidogenic proteins including amyloid-β, tau, and superoxide dismutase 1 G93A – key biomarkers in Alzheimer’s and amyotrophic lateral sclerosis (ALS) disease pathologies.^28,29^ Further evidence for a protective role comes from FAIM knockout models, where loss of FAIM results in increased protein aggregation and susceptibility to oxidative and thermal stress.^30^ In the metabolic disease space, one study proposed that FAIM contributes to the regulation of insulin signaling by regulating the expression of IRβ and IRS and activation of Akt.^31^ While these findings suggest that FAIM may contribute to both proteostasis and metabolic regulation, there are currently no reports demonstrating a role for FAIM in the aggregation of insulin or other endocrine proteins.

Given FAIM’s ubiquitous expression and activity against aggregation-prone proteins, we hypothesized that FAIM may also regulate insulin aggregation. To test this possibility, we examined the effects of FAIM-S on insulin fibrillation using complementary biophysical, structural, and cellular approaches. Aggregation kinetics were monitored by ThT fluorescence, fibril morphology was evaluated by transmission electron microscopy (TEM), and structural changes were characterized by circular dichroism (CD) and Fourier transform infrared (FTIR) spectroscopy. Molecular level interactions were further investigated using solution-state ^1^H NMR spectroscopy, while aggregate associated cytotoxicity was assessed using cell viability assays. Our findings demonstrate that FAIM modulates insulin aggregation, alters fibril morphology, reduces aggregate associated toxicity, and interacts with regions of insulin implicated in amyloid formation. These results identify FAIM as a regulator of insulin aggregation and expand the growing repertoire of endogenous mechanisms that suppress amyloid formation.

## Results

### FAIM attenuates insulin fibril formation

To investigate whether FAIM modulates insulin fibrillation, we monitored amyloid formation by ThT fluorescence at pH 3.0 (Fig. 1). Reactions containing 80 μM insulin were incubated at 37 °C under continuous shaking in the absence or presence of 5, 10, or 20 μM FAIM. Insulin alone exhibited sigmoidal fluorescence kinetics with a lag phase of approximately 24 hours (Fig. 1a). These kinetics are consistent with nucleation-dependent fibrillation of insulin reported under similar acidic conditions.^32^ FAIM reduced ThT fluorescence at all concentrations tested, indicating decreased formation of ThT-positive aggregates. The greatest reduction was observed at 5 μM FAIM, whereas 10 and 20 μM FAIM produced progressively higher fluorescence signals, although both remained substantially lower than those of insulin alone (Fig. 1b). Notably, samples containing 20 μM FAIM exhibited a gradual increase in fluorescence throughout the experiment. These observations agreed with previous reports showing that chaperone efficacy often depends on stoichiometry.^33–36^ For example, sub-stoichiometric inhibition as observed with DNAJB6 for Aβ42 can be highly effective, but the exact ratio is protein- and context-dependent.^36^ Accordingly, we sought to determine whether the greater inhibition observed at lower FAIM concentrations reflected a favorable FAIM-to-insulin ratio by repeating the assay at a fixed FAIM molar ratio of 1:5 across multiple protein concentrations (Fig. S1). FAIM suppressed insulin aggregation under all conditions tested, and the lowest FAIM concentration again produced the greatest reduction in ThT fluorescence. This observation suggests that the concentration dependence of FAIM-mediated inhibition is not solely a consequence of the FAIM-to-insulin ratio. Rather, factors in addition to chaperone stoichiometry likely contribute to the inhibitory activity of FAIM.

**Fig. 1.**
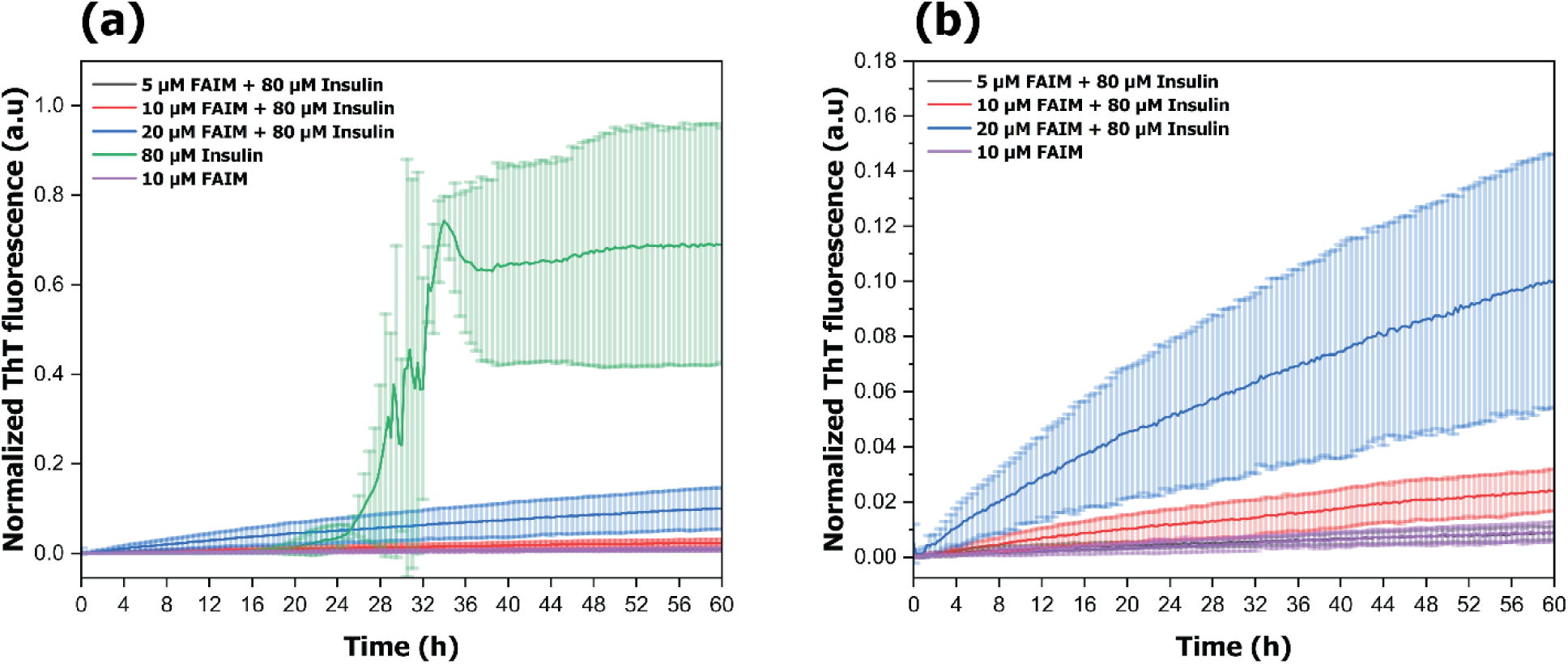
**(a)** ThT fluorescence kinetics of 80 μM insulin incubated in the absence or presence of 5, 10, or 20 μM FAIM. FAIM alone (10 μM) is included as a control. **(b)** Expanded view of the FAIM-containing samples shown in **(a)**. Conditions: 10 mM sodium phosphate, 150 mM NaCl, 10 μM ThT, pH 3.0, 37 °C, 700 rpm. Data represent mean ± SD from three independent experiments.

### FAIM alters insulin fibril morphology and structure

To determine whether FAIM altered the morphology of insulin aggregates, samples were examined by transmission electron microscopy (TEM) following 48 h of incubation (Fig. 2). Insulin alone formed dense networks of long unbranched fibrils typical of mature amyloid aggregates.^37,38^ Fibrils remained visible at 5 µM FAIM but appeared thinner and less entangled than those formed by insulin alone (Fig. 2a). At 10 µM FAIM, fibrillar structures were less abundant and frequently appeared fragmented. Increasing the concentration to 20 µM resulted in a near-complete loss of fibrillar morphology. Instead, the sample contained irregular nonfibrillar aggregates lacking obvious long-range order (Fig. 2c). The TEM data revealed a more complex relationship between aggregate morphology and ThT fluorescence. Although 5 μM FAIM produced the greatest reduction in ThT fluorescence, fibrillar structures remained evident by TEM. By contrast, samples containing 20 μM FAIM exhibited both reduced ThT fluorescence and a substantial loss of recognizable fibrils. It is therefore possible that the reduced ThT fluorescence does not reflect a proportional reduction in aggregate abundance and may instead arise from FAIM-induced changes in aggregate structure. These findings warrant further characterization of the aggregates formed in the presence of FAIM.

**Fig. 2.**
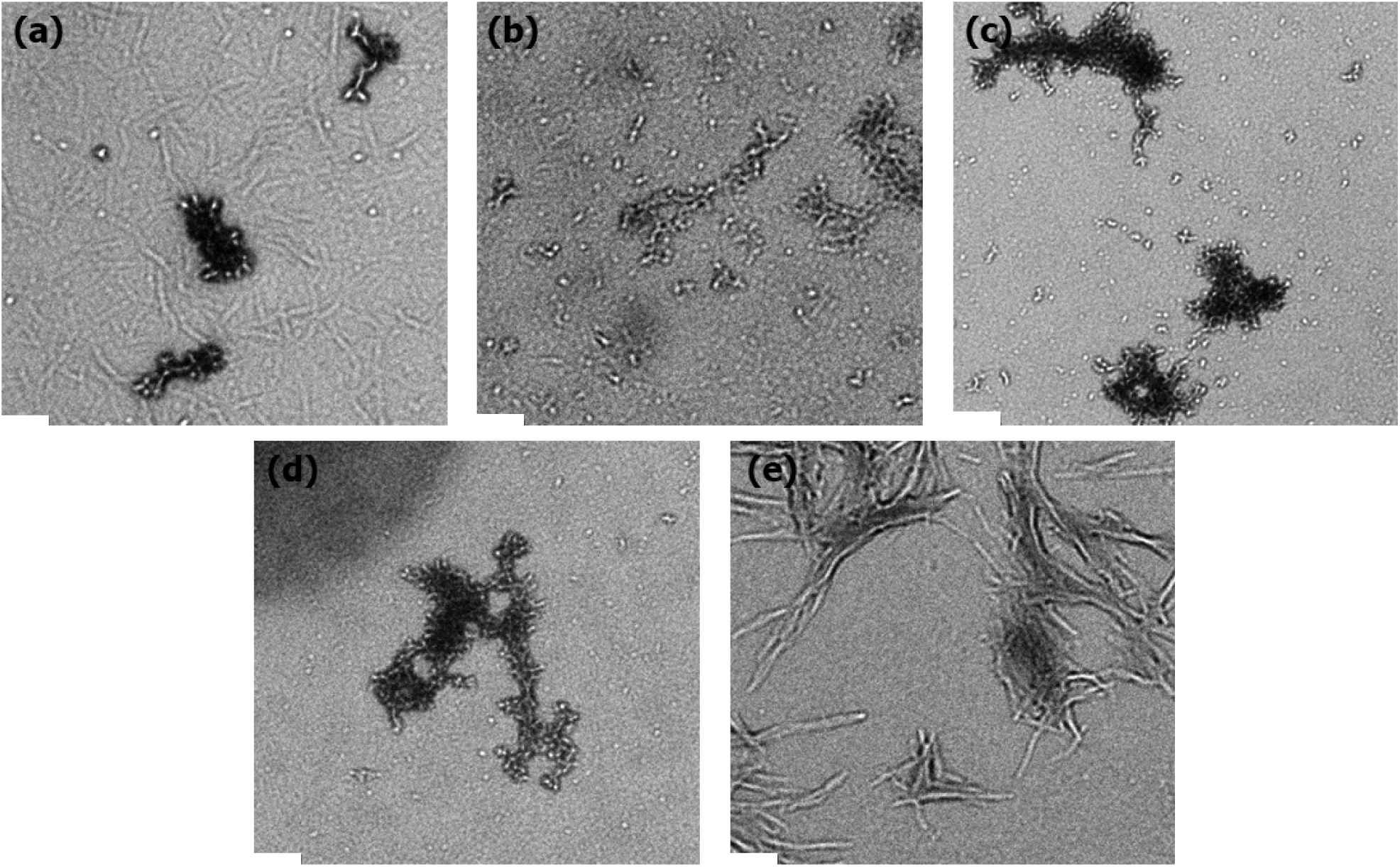
TEM images acquired after 48 h of incubation of insulin with (a) 5 µM FAIM, (b) 10 µM FAIM, and (c) 20 µM FAIM. Control samples containing (d) FAIM alone and (e) insulin alone are also shown. Samples were incubated at 37 °C with agitation (700 rpm) in sodium phosphate buffer (pH 3.0), negatively stained with uranyl acetate, and imaged by TEM. Representative images from three independent sample preparations are shown. Scale bar, 200 nm.

While TEM revealed substantial changes in aggregate morphology, it could not determine whether these differences were accompanied by alterations in secondary structure. To examine this possibility, Fourier-transform infrared (FTIR) spectra were collected in the amide I region (1550–1800 cm□^1^), which is sensitive to protein backbone conformation (Fig. 3; Fig. S2).^39^ Insulin aggregates formed in the absence of FAIM exhibited a dominant peak at ∼1628 cm□^1^, commonly assigned to β-sheet-rich structures.^40,41^ The spectra changed progressively upon addition of FAIM. At 10 µM and 20 µM FAIM, the amide I peak shifted modestly toward higher wavenumbers relative to insulin alone. A much larger shift was observed at 5 µM FAIM, where the dominant peak occurred near 1655 cm□^1^. The position of this peak is generally associated with α-helical or disordered conformations. The attenuation and shift of the amide I peak mark a transition from β-sheet–rich structures to more disordered or alternative aggregate forms.^42^

**Fig. 3.**
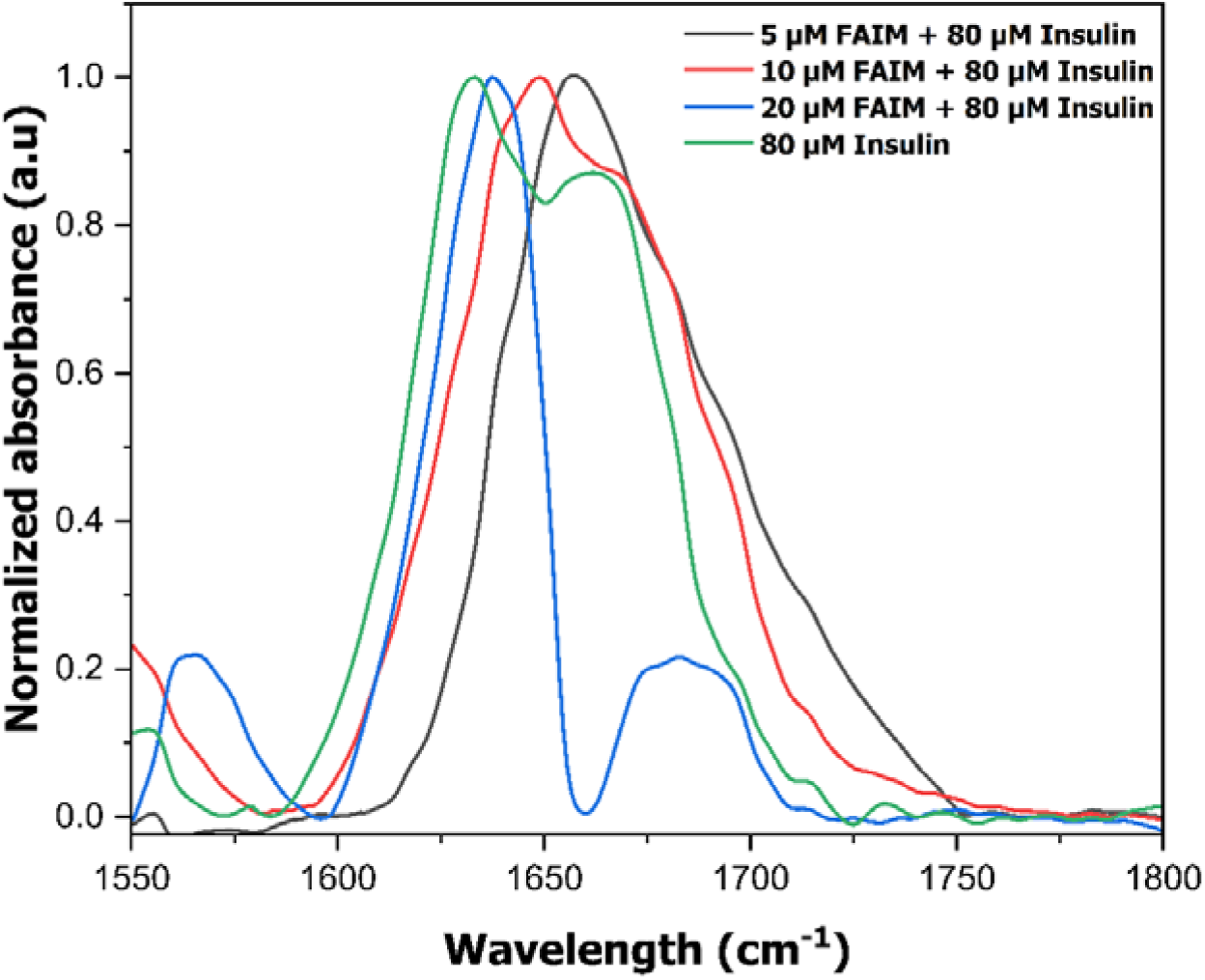
FTIR spectra in the amide I region (1550-1800 cm□^1^) of aggregates formed from 80 µM insulin incubated alone or in the presence of 5, 10, or 20 µM FAIM for 48 h at 37 °C with continuous agitation. Corresponding Gaussian deconvolutions used for secondary structure analysis are shown in Fig. S2.

Far-UV circular dichroism (CD) spectroscopy corroborated the structural changes observed by FTIR (Fig. 4). Our group has previously reported insulin in its native monomeric state is predominantly α-helical, with characteristic double minima near 210 and 222 nm.^43^ Conversely, insulin aggregates formed in the absence of FAIM displayed a broad minimum centered near 230 nm, confirming a β-sheet dominated conformation as observed in amyloids.^44,45^ The addition of FAIM altered this profile at all concentrations tested. Rather than the broad minimum observed for insulin alone, samples containing FAIM exhibited spectra with minima near 210 and 222 nm, indicating increased α-helical character and reduced β-sheet content.^46^ The CD spectrum of FAIM alone showed minima near 218 nm which is consistent with β-sheet structure. Because CD intensities reflect ensemble contributions from both proteins, their apparent non-linearity with FAIM concentration likely stems from overlapping signals of insulin and FAIM. FTIR analysis resolved this trend more clearly. 5 µM FAIM produced the strongest shift to a helical structure, 10 µM retained partial helix, and 20 µM reverted toward greater β-sheet content. Secondary structure estimates from BeStSel analysis supported this interpretation and identified the greatest increase in helical content together with the largest reduction in β-sheet structure at 5 µM FAIM (Table S1). The agreement between the morphological and spectroscopic measurements suggests that FAIM alters the structural organization of insulin aggregates. Rather than forming the β-sheet-rich fibrils observed for insulin alone, insulin incubated with FAIM gave rise to aggregate populations with distinct structural properties.

**Fig. 4.**
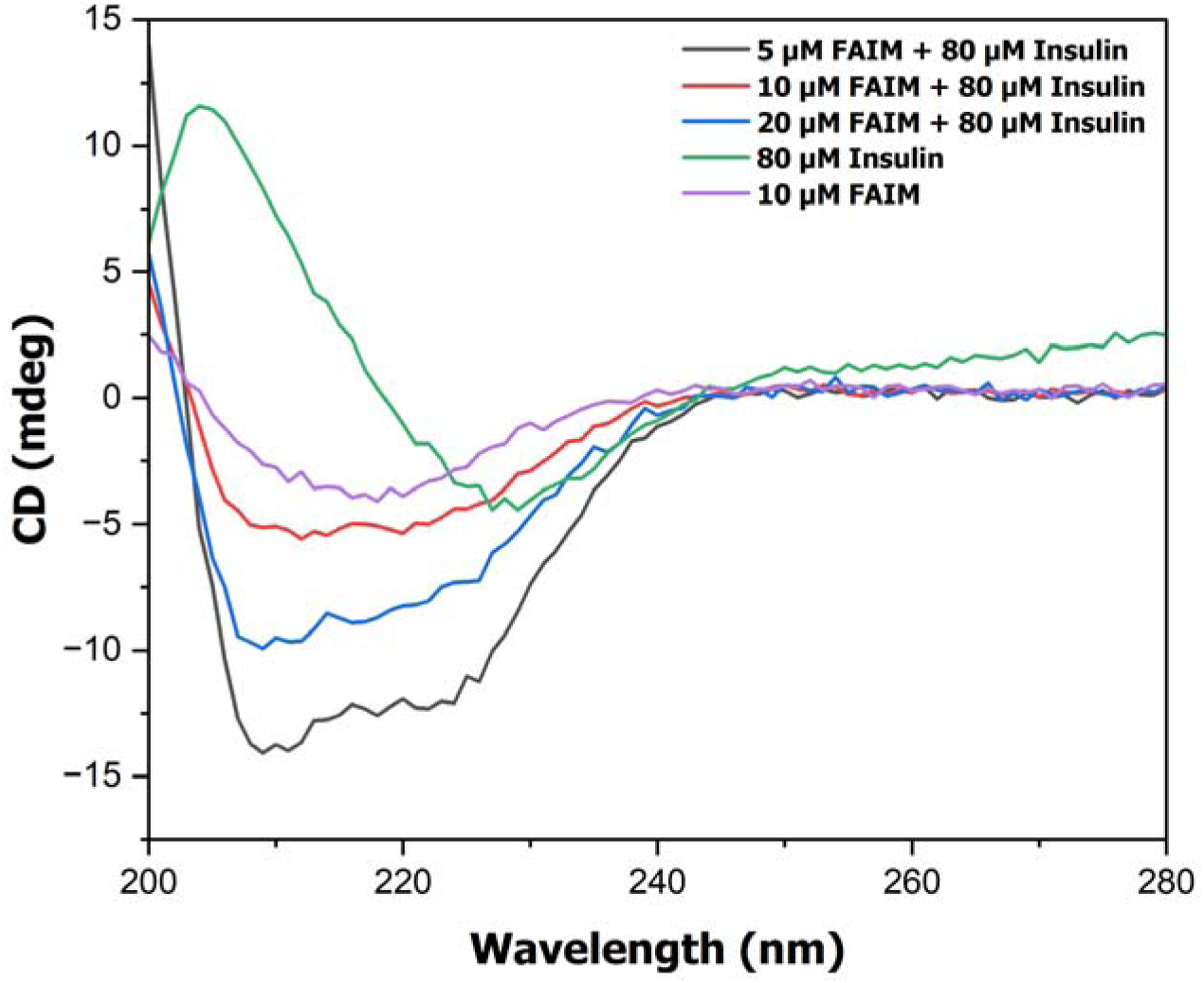
Far-UV CD spectra (200-280 nm) of 80 µM insulin incubated alone or with 5, 10, or 20 µM FAIM for 48 h at 37 °C under continuous agitation. A spectrum of 10 µM FAIM alone is included as a control. Quantitative secondary structure estimates derived from spectral deconvolution are provided in Table S1.

### FAIM reduces insulin aggregate induced cytotoxicity

To assess whether FAIM-mediated changes in insulin aggregation altered aggregate cytotoxicity, we performed an XTT cell viability assay using a NIH3T3 mouse fibroblast line (Fig. 5). Insulin (80 μM) was aggregated for 48 h alone or in the presence of 5 or 20 μM FAIM before dilution to final insulin concentrations of 5, 10, 20, or 40 μM for cellular studies. Insulin aggregates reduced cell viability at all concentrations tested. Aggregates formed in the presence of FAIM were consistently less toxic than those formed by insulin alone. The greatest protection was observed for aggregates generated with 5 μM FAIM, which significantly improved viability at final insulin concentrations of 5, 10, and 20 μM (p = 0.0001, 0.0167, and 0.0058, respectively). Aggregates formed with 20 μM FAIM produced a smaller effect and reached statistical significance only at the lowest insulin concentration tested (p = 0.0006). At 40 μM insulin, viability was comparable across all treatment groups. The concentration dependence observed in the cell viability assay paralleled that observed in the biophysical experiments, with the largest effects consistently occurring in the presence of 5 μM FAIM. The reduced cytotoxicity coincided with decreased ThT fluorescence, altered aggregate morphology, and changes in aggregate secondary structure, suggesting that FAIM redirects insulin aggregation toward structurally distinct assemblies with diminished toxic potential. It is important to note however because we did not directly characterize oligomeric species or mechanisms of cell damage, we cannot conclude which specific aggregate populations are responsible for the observed differences in viability.

**Fig. 5.**
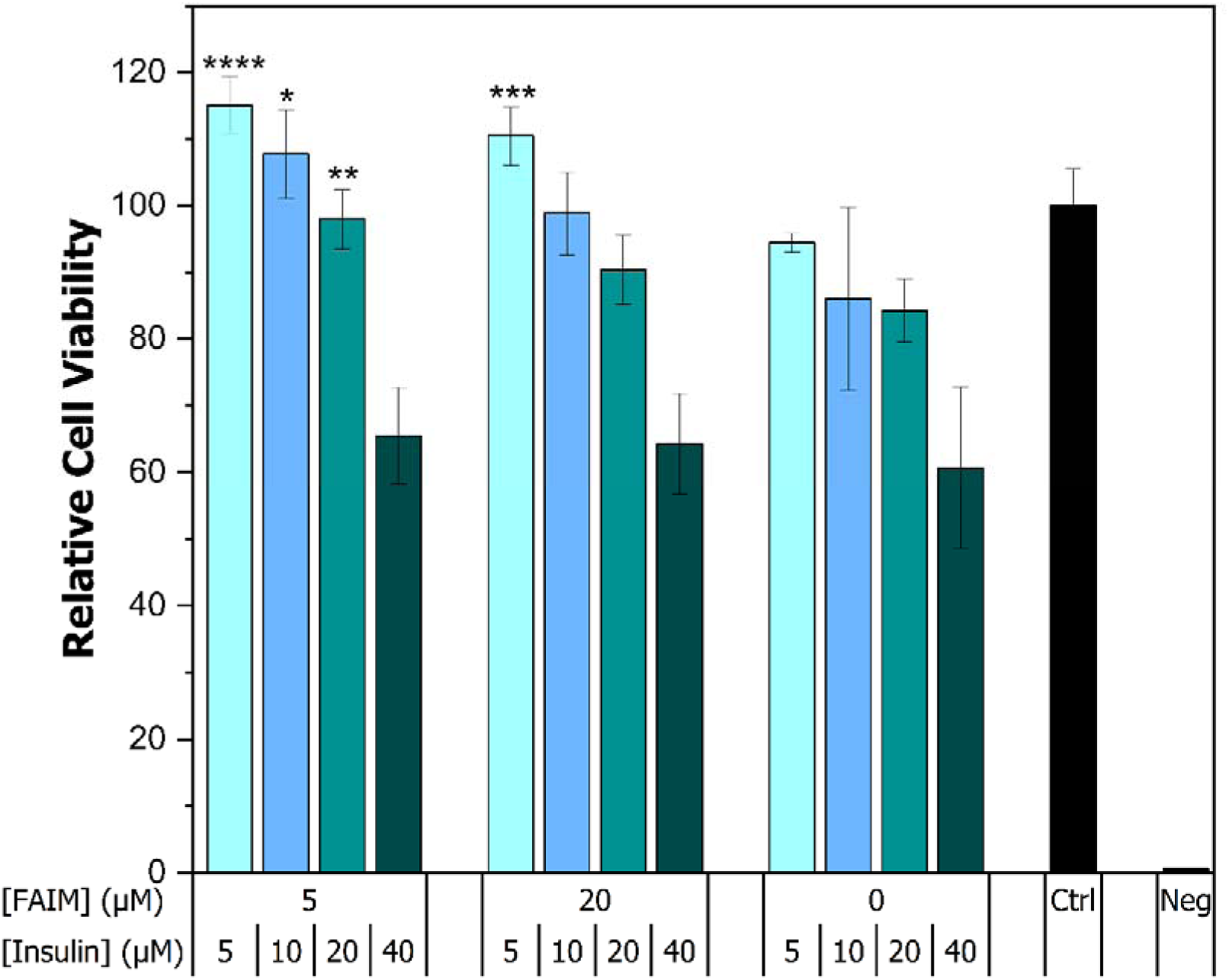
Relative viability of NIH3T3 fibroblasts following exposure to preformed insulin aggregates generated in the absence or presence of 5 or 20 µM FAIM. Aggregates were added to cells at final insulin concentrations of 5, 10, 20, or 40 µM. Untreated cells served as the 100% viability control, and 0% isopropanol was included as a negative control. Cell viability was assessed after 24 h using an XTT assay. Data represent mean ± SD from four independently treated wells per condition. Statistical significance is shown relative to insulin alone at the corresponding insulin concentration (*p < 0.05, **p < 0.01, ***p < 0.001, ****p ≤ 0.0001).

### FAIM preserves insulin solubility and molecular mobility during aggregation

^1^H NMR spectroscopy was employed to investigate the effects of FAIM on the solution-state behavior of insulin throughout the aggregation process (Fig. 6). A sample of 10 µM FAIM and 80 µM insulin were prepared under acidic conditions conducive to fibril formation, and spectra were acquired at 0, 6, 12, 24, and 48 hours at 37 °C. In the absence of FAIM, insulin resonances progressively decreased in intensity over time, particularly within the amide/aromatic (6-9 ppm) and aliphatic (0.5-2 ppm) regions. By 24 h, most signals had disappeared into the baseline, consistent with the conversion of soluble insulin into large aggregates that are not detected by solution-state NMR because of their slow molecular tumbling.^47,48^ A similar loss of signal intensity was observed in samples containing FAIM, but the effect occurred substantially earlier. Most resonances were markedly attenuated by 6 h and remained absent throughout the remainder of the experiment. The accelerated disappearance of NMR signals indicates a more rapid depletion of NMR-visible insulin species in the presence of FAIM. Together with our other data, these observations suggest that FAIM alters the aggregation pathway of insulin, leading to the formation of aggregate species that differ from those formed by insulin alone. Comparison of the 0 h spectra provided insight into the initial stages of the interaction (Fig. 7). The spectra of insulin alone and insulin incubated with FAIM were nearly superimposable, with no detectable chemical shift perturbations or peak broadening. Although FAIM exhibited a distinct set of resonances when analyzed alone, its addition did not measurably alter the insulin spectrum. These results indicate that FAIM does not induce a detectable structural perturbation of insulin immediately upon mixing. Any interactions present at the start of the reaction are therefore likely to be transient, weak, or involve only a small fraction of the insulin population. The NMR data therefore suggests that the effects of FAIM emerge during the aggregation process rather than through a stable interaction with monomeric insulin.

**Fig. 6.**
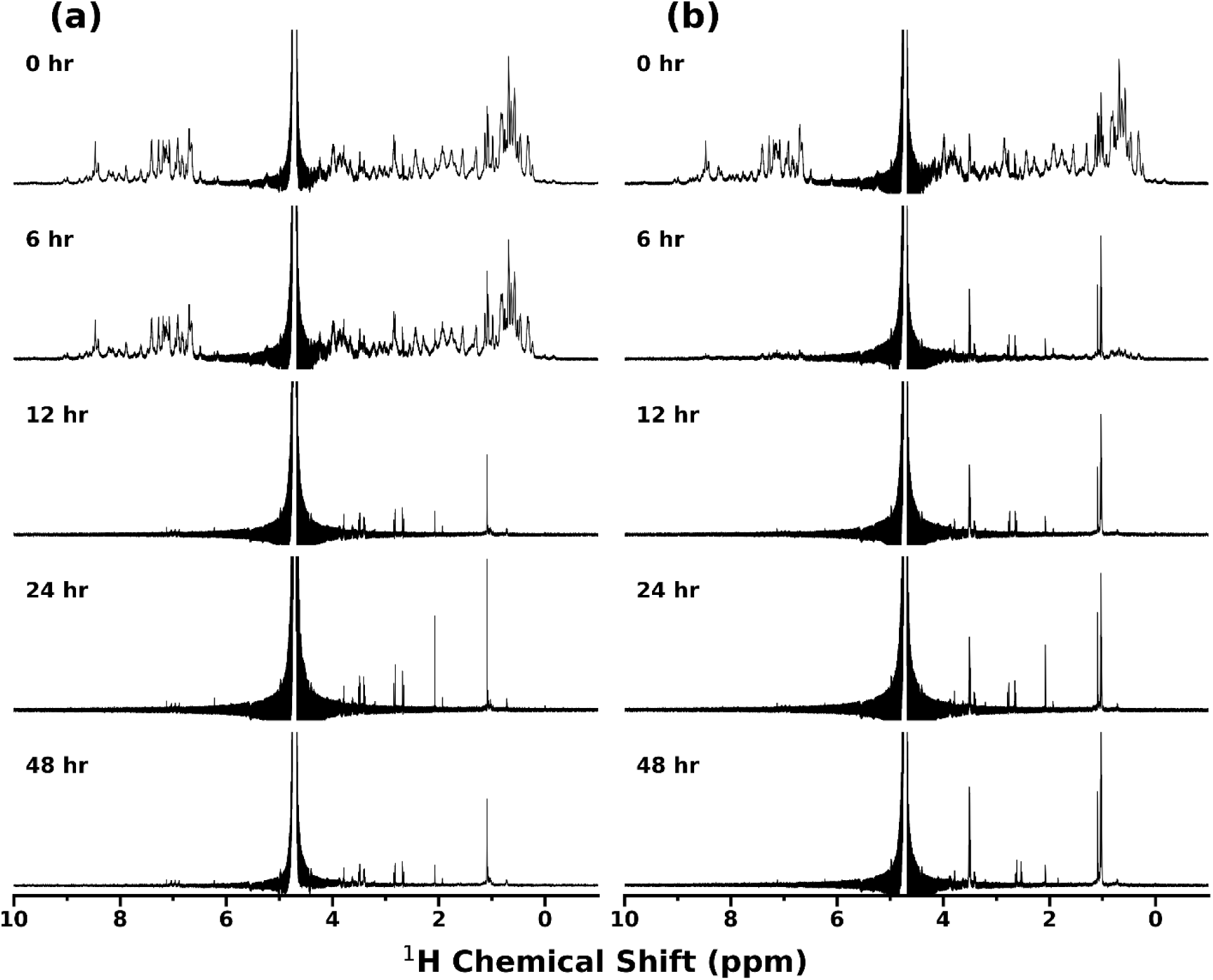
Time-resolved ^1^H NMR spectra of **(a)** 80 µM insulin incubated alone and **(b)** 80 µM insulin incubated with 10 µM FAIM. Spectra were acquired at 0, 6, 12, 24, and 48 h during aggregation in 10 mM sodium phosphate buffer (pH 3.0) at 37 °C with continuous agitation (700 rpm).

**Fig. 7.**
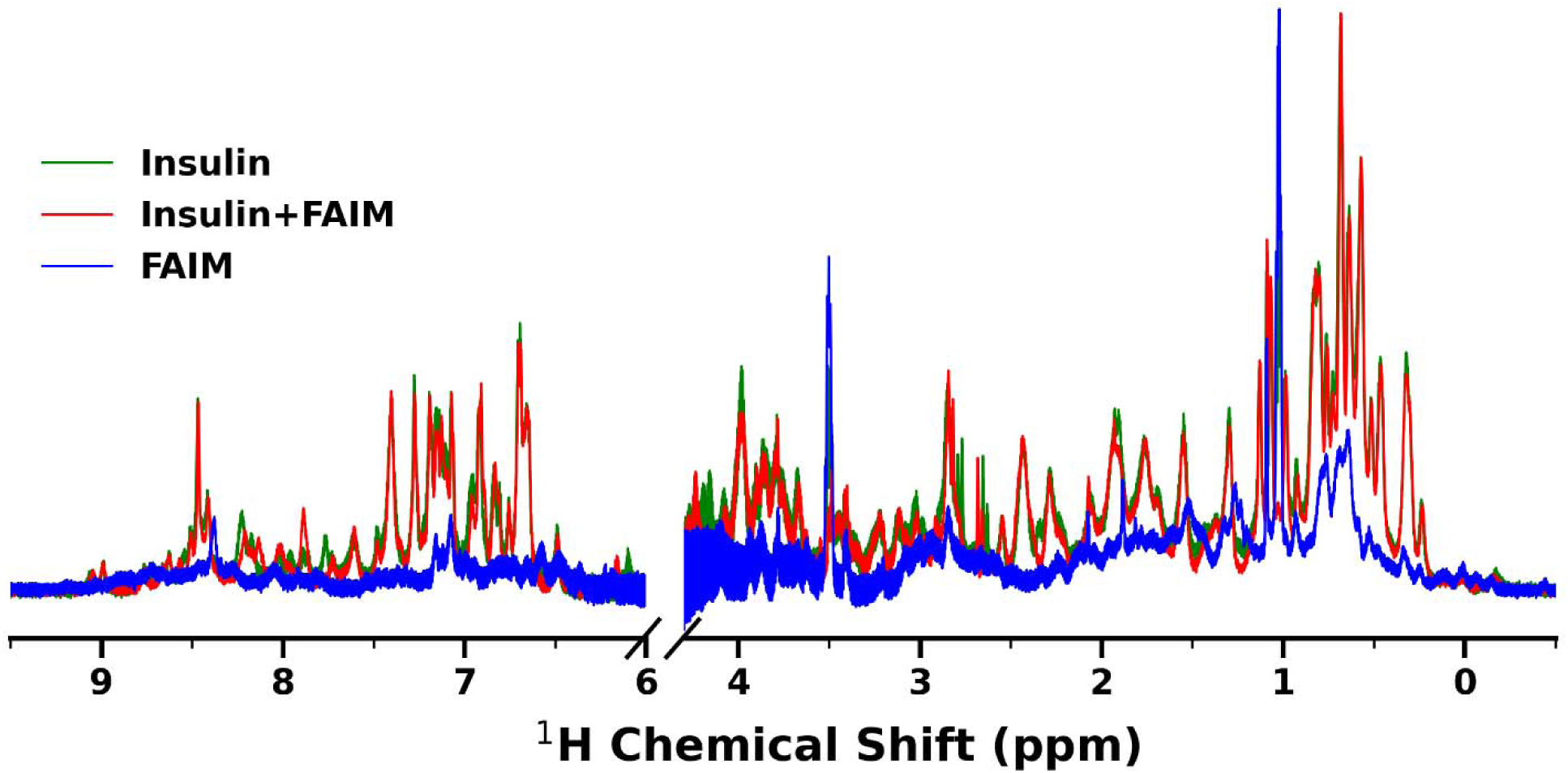
Overlay of ^1^H NMR spectra of 10 µM FAIM, 80 µM insulin, and a mixture containing 80 µM insulin with 10 µM FAIM at 0-hour time point in 10 mM sodium phosphate buffer (pH 3.0, 10% D O) at 37 °C. Spectra span the aliphatic, aromatic, and amide proton regions (0-9.4 ppm).

### Predicted structure of the FAIM-insulin complex at pH 3.0

Structural modeling predicted that FAIM engages a composite binding surface spanning both insulin chains at the A/B-chain interface (Fig. 8). Pairwise heavy-atom distance analysis identified 67 residue pairs within contact distance, with the closest interactions occurring at approximately 2.5-3.0 Å. FAIM contributed basic and polar residues (Lys110, Lys130, Lys175, Arg174, His180, Ser170, Thr153, Asp150) that clustered at the interface. These residues contacted A-chain Glu17, Asn18, and Asn21, and B-chain Asn3, His10, Glu13, Leu17, Glu21, Arg22, and Pro28. Interactions included Lys130–A:Glu17, Arg174–B:Glu21, Glu176–B:Arg22, and Asp150–B:His10. Although these resemble salt bridges in neutral conditions, at pH 3.0 the acidic groups were protonated; they are therefore best described as electrostatic contacts. Additional hydrogen bond-like interactions were observed between FAIM Lys110/Lys175 and insulin Asn18/Asn21, and between FAIM Ser170/Thr153 and insulin Asn3/Arg22. Hydrophobic and van der Waals contacts, particularly between FAIM Gly172/Lys103/Lys110 and B-chain Leu17/Pro28, further stabilized the interface. Interestingly, the disulfide bonds within insulin (three SG–SG linkages detected between A- and B-chain cysteines) remained intact in the complex, indicating that FAIM binding does not disrupt the native insulin fold. These results describe FAIM associating with insulin via a network of complementary electrostatic, polar, and hydrophobic contacts, targeting residues at the junction of the A and B chains while preserving insulin’s disulfide structure. This model provides a mechanistic basis for the inhibitory activity observed in our biophysical characterization experiments. By clustering around aggregation-prone hotspots (A:Glu17, A:Asn18, A:Asn21, B:Glu21, B:Arg22), FAIM masks surfaces that normally drive β-sheet assembly. By stabilizing these regions through electrostatic and hydrogen bond–like contacts, FAIM likely disrupts the earliest steps of fibril formation, prolonging the lag phase and reducing fibrils at low concentrations. The contribution of hydrophobic contacts at B:Leu17 and B:Pro28 further blocks surfaces needed for fibril elongation. By selectively binding to key nucleation residues, relatively small amounts of FAIM are sufficient to alter early assembly events. However, the precise molecular determinants of FAIM-mediated inhibition remain to be established experimentally.

**Fig. 8.**
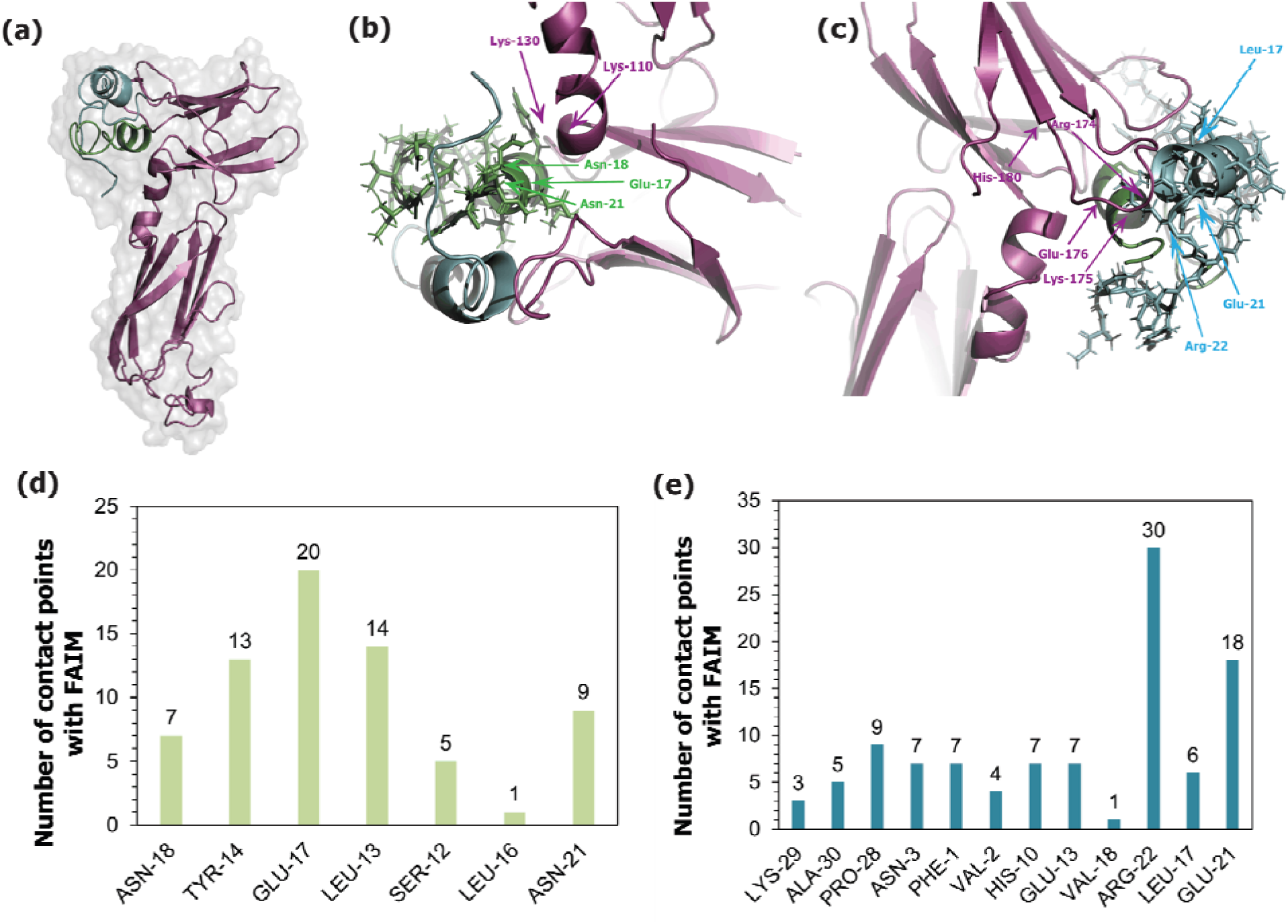
PyMOL representations of the predicted FAIM-insulin complex at pH 3.0. **(a)** Overall structure of the complex, with FAIM shown in purple, the insulin A-chain in green, and the insulin B-chain in teal. Protein surfaces are rendered semi-transparent to visualize the interaction interface. **(b)** Expanded view of the insulin A-chain interface highlighting contacts with residues Glu17, Asn18, and Asn21. **(c)** Expanded view of the insulin B-chain interface showing interactions involving residues Glu21, Arg22, and Leu17. **(d,e)** Residue contact analysis of the insulin A- and B-chains, respectively. Bar plots show the number of heavy-atom contacts formed with FAIM.

## Discussion

Insulin aggregation has been extensively characterized under fibril-promoting conditions *in vitro*,^49–54^ yet less is known about the endogenous factors that may influence this process. To this end, we investigated FAIM, a stress-responsive protein implicated in proteostasis, and studied its effect on insulin fibrillation. A clear inhibitory effect was observed, with FAIM reducing ThT fluorescence at all concentrations tested. Notably, the greatest inhibition occurred at 5 µM FAIM despite insulin being present in substantial molar excess. Although this finding indicates that FAIM effectively perturbs the aggregation process, the underlying mechanism was not immediately apparent. As discussed below, structural, spectroscopic, and cellular analyses revealed that the effects of FAIM extend beyond a simple reduction in fibril formation and point instead to changes in the nature of the aggregates that are formed.

The concentration dependence of the inhibitory effect was unexpected. Although FAIM reduced ThT fluorescence at all concentrations tested, the greatest inhibition was consistently observed at 5 µM FAIM, with progressively higher fluorescence signals observed at 10 and 20 µM. This trend was also observed at a fixed FAIM-to-insulin ratio across multiple protein concentrations, indicating that the enhanced inhibition at lower FAIM concentrations cannot be explained solely by the stoichiometric relationship between FAIM and insulin. The pronounced effect observed at 5 µM FAIM is noteworthy because insulin was present in a 16-fold molar excess, suggesting that relatively small amounts of FAIM are sufficient to perturb the aggregation process. Consistent with previous reports that small chaperone proteins can modulate amyloid formation through interactions with aggregation intermediates,^55,56^ similar sub-stoichiometric effects have been observed for other inhibitors of amyloid assembly. For example, Hsp70 inhibits IAPP aggregation at sub-stoichiometric concentrations through preferential interactions with low abundance prenucleation oligomers while exhibiting relatively weak binding to monomeric protein.^57^ Likewise, the mitochondrial-derived peptide HumaninS14G suppresses IAPP aggregation at molar ratios as low as 1:250, leading the authors to conclude that inhibition does not require bulk capture of monomeric IAPP.^58^ Together, these studies suggest that effective suppression of amyloid formation can be achieved through interactions with a limited population of aggregation-prone species. Whether a similar mechanism underlies FAIM-mediated inhibition remains uncertain. Because 5 µM represented the lowest FAIM concentration examined, we cannot exclude the possibility that maximal inhibition occurs at concentrations below the range tested in this study.

Our NMR data provides additional insight into this possibility. No detectable changes in the insulin spectrum were observed immediately following the addition of FAIM, indicating that FAIM does not measurably perturb the native monomer prior to aggregation. While weak or transient interactions cannot be excluded, the absence of chemical shift perturbations or peak broadening suggests that any interaction with monomeric insulin is limited under these conditions. Our results are consistent with a number of previous studies showing that molecular chaperones, especially ATP-independent sHsps and related co-chaperones, preferentially engage aggregation-prone conformers or oligomeric intermediates with relatively weak affinity for native proteins.^59–61^ This selective binding serves to trap non-native species, prevent their further aggregation, and channel them toward productive refolding or degradation pathways as part of a dynamic proteostasis network. However, the present data does not distinguish whether these species correspond to partially unfolded conformers, oligomeric intermediates, or early aggregates.

An important finding related to the molecular structure of the aggregates formed in the presence of FAIM. The TEM, CD, and FTIR analyses pointed to changes in aggregate morphology and secondary structure that depended on the FAIM concentration. At first glance, these observations may seem difficult to reconcile with the fluorescence data. Despite producing the greatest reduction in ThT fluorescence, 5 µM FAIM did not eliminate fibril formation, as fibrillar structures remained detectable by TEM. This could reflect the formation of structurally distinct aggregate species rather than a simple reduction in aggregate formation. Thus, the persistence of fibrillar material does not necessarily imply the formation of conventional insulin amyloid fibrils.

Further support for this interpretation was provided by the CD and FTIR measurements. At 5 µM FAIM, CD spectra revealed increased α-helical character accompanied by reduced β-sheet content, while FTIR analysis showed a shift of the amide I band toward higher wavenumbers consistent with diminished cross-β structure. These observations provide a potential explanation for the reduced ThT fluorescence despite fibrillar material observed by TEM. Specifically, the aggregates formed in the presence of FAIM may retain fibrillar morphology while possessing altered secondary structural organization that limits ThT binding. The mechanism underlying this effect remains unclear. Although FAIM itself exhibits β-sheet character, our data does not suggest that it promotes β-sheet formation in insulin. Instead, FAIM may interfere with the alignment or packing of β-strands required for the development of mature amyloid architecture. The increased α-helical signal observed in the presence of FAIM agrees with prior studies describing aggregation modulators that stabilize helical intermediates or molten-globule-like states during protein misfolding.^62,63^ Yet it is worth noting that we cannot determine from the present data whether these structural changes arise from direct incorporation of FAIM into insulin aggregates or from transient interactions with aggregation intermediates.

Interestingly, the structural changes induced by FAIM were accompanied by reduced aggregate-associated cytotoxicity. NIH3T3 fibroblasts exposed to aggregates formed in the presence of 5 µM FAIM exhibited greater viability than cells treated with aggregates generated from insulin alone, whereas aggregates formed in the presence of 20 µM FAIM were only modestly protective. This trend parallels the concentration dependence observed in the ThT experiments, where the greatest inhibition was likewise observed at 5 µM FAIM. A previous inhibitor study reported a similar observation in the context of huntingtin exon 1 aggregation.^64^ Sub-stoichiometric concentrations of curcumin altered aggregate structure and morphology, and the resulting aggregates induced significantly less cellular stress than those formed in the absence of inhibitor. Taken together, these observations suggest that the structural remodeling induced by FAIM is associated with a reduction in aggregate toxicity.

While we do not have direct experimental evidence defining the interaction interface between FAIM and insulin, the predicted complex provides a possible framework for interpreting the observed inhibitory activity. In the model, FAIM engages residues located near the insulin A/B-chain interface, including several residues that have previously been implicated in insulin self-association and amyloid formation.^15,65^ This observation was significant given low concentrations of FAIM were sufficient to alter aggregation structure and reduce cytotoxicity. Previous structural and mutational studies have shown that localized perturbations within regions that govern oligomerization can substantially alter aggregation behavior without requiring global disruption of the protein structure.^66,67^ Such findings suggest that relatively limited interactions can have disproportionately large effects on amyloid assembly by shifting the conformational landscape away from productive nucleation and elongation pathways.

In this context, the predicted interaction of FAIM with residues involved in insulin self-association may provide a structural basis for the effects observed at sub-stoichiometric concentrations. Rather than preventing aggregation through extensive binding to monomeric insulin, FAIM may perturb a small number of critical intermolecular interactions that influence the subsequent course of assembly. At the same time, the relationship between the predicted complex and the species present during aggregation remains uncertain. The modeled contacts may reflect interactions with partially unfolded conformers or early aggregates rather than native insulin. As such, the docking model provides a useful structural hypothesis, but does not establish the precise molecular mechanism by which FAIM alters insulin aggregation.

In summary, our results suggest that FAIM functions as an endogenous modulator of insulin aggregation by altering the structural and toxic properties of insulin assemblies formed under fibril promoting conditions. We propose that FAIM interacts with aggregation-prone insulin species and redirects the aggregation pathway toward alternative assemblies with reduced cytotoxicity. Our study focused on insulin fibrillation at acidic pH, although the possibility remains that FAIM may exert similar effects under other aggregation-promoting conditions. Future work may define the structural determinants governing this interaction and establish whether related mechanisms influence the aggregation of other amyloidogenic proteins. Collectively, our findings demonstrate that relatively low concentrations of FAIM can substantially alter both the structural and biological properties of insulin aggregates and suggest that endogenous protein quality-control factors may influence the outcome of amyloid formation in ways that extend beyond simple suppression of aggregation.

## Materials and methods

### Materials

Recombinant unlabeled FAIM-S was expressed and purified as previously described by Dr. Rothstein’s laboratory at Western Michigan University Homer Stryker M.D. School of Medicine (Kalamazoo, MI, USA). Recombinant human insulin was obtained from Roche (Indianapolis, IN, USA). ThT dye and TEM copper grids were purchased from MilliporeSigma (Burlington, MA, USA). Sodium chloride, sodium phosphate, and trypsin–EDTA were purchased from Fisher Scientific (Hampton, NH, USA). UranyLess stain was obtained from Electron Microscopy Sciences (Hatfield, PA, USA). Dulbecco’s Modified Eagle Medium (DMEM) was purchased from Gibco (Grand Island, NY, USA).

### ThT fluorescence assay

Fresh insulin stock solutions were prepared in 10 mM sodium phosphate buffer (pH 3.0), and concentrations were determined spectrophotometrically using a NanoPhotometer NP80 (IMPLEN) and an extinction coefficient of 3,360 M□^1^ cm□^1^. FAIM-S stock solutions were prepared in 10 mM sodium phosphate buffer (pH 7.4), and concentrations were determined using an extinction coefficient of 32,555 M□^1^ cm□^1^. Samples were supplemented with 10 μM ThT and 150 mM NaCl and loaded (50 μL per well) in triplicate into black walled, flat bottom 384-well microplates (Costar, 0976121). Fluorescence measurements were acquired using a Synergy H1 multimode microplate reader (Agilent BioTek) maintained at 37 °C with continuous orbital shaking (700 rpm) for 48 h. ThT fluorescence was monitored using excitation and emission wavelengths of 452 and 485 nm, respectively. Measurements were collected at 15-minute intervals. Fluorescence values were normalized to baseline readings and averaged across triplicates for each concentration. Error bars represent one standard deviation from three independent experiments. Data plots and statistical analyses were completed using OriginPro 2025 (OriginLab).

### TEM

Aggregate morphology was assessed by TEM following completion of the ThT fluorescence assays. Aliquots (10 μL) of each sample were deposited onto hydrophilic 300-mesh carbon-coated Formvar copper grids (MilliporeSigma, TEM-FCF300CU) and allowed to adsorb for 10 minutes at room temperature. Grids were then negatively stained with 10 μL uranyl acetate (Electron Microscopy Sciences, 22409) and air dried at room temperature before imaging. Images were acquired using an HT7800 transmission electron microscope (Hitachi) operated at an accelerating voltage of 100 kV. Micrographs were collected from at least three independent regions of each grid at magnifications ranging from 15,000x to 25,000x.

### CD spectroscopy

After 48 hours of incubation, CD samples were prepared by diluting 30 μL aliquots (80 μM insulin containing 5, 10, or 20 μM FAIM-S) or control solutions (80 μM insulin and 10 μM FAIM-S) into 170 μL of 10 mM sodium phosphate buffer (pH 3.0). CD spectra were acquired using a Chirascan Plus spectropolarimeter (Applied Photophysics). For each sample, spectra were recorded as the average of three accumulations. Measurements were collected using a scanning speed of 100 nm min□^1^, bandwidth of 1 nm, data pitch of 0.5 nm, integration time of 1 s, and CD scale of 200 mdeg. Processed spectra were plotted using OriginPro 2025 (OriginLab) and analyzed in BestSel (v1.3.230210) to evaluate secondary structure. Deconvolution was performed by uploading baseline-corrected spectra in mdeg, with a specified pathlength of 0.1 cm and a protein concentration of 80 μM.

### FTIR

FTIR spectra were acquired using a Thermo Nicolet spectrometer equipped with a diamond attenuated total reflectance (ATR) accessory (Thermo Fisher Scientific, Waltham, MA, USA). For each sample, 512 scans were averaged across the spectral range of 1500-1800 cm□^1^ at a resolution of 8 cm□^1^. Spectra were subjected to baseline correction and spectral deconvolution of the amide I region (1600-1700 cm□^1^) using OriginPro 2025 (OriginLab). Component bands were fitted using mixed Gaussian-Lorentzian functions, with initial peak positions determined from second-derivative spectra. Peak assignments were based on established amide I band frequencies corresponding to β-sheet, α-helical, turn, and random-coil secondary structures.

### Cellular cytotoxicity

NIH3T3 mouse fibroblasts were cultured in Dulbecco’s modified Eagle medium (DMEM) supplemented with 10% fetal bovine serum and 1% penicillin-streptomycin at 37 °C in a humidified atmosphere containing 5% CO□. Cells were passaged at 70-80% confluency using 0.25% trypsin-EDTA and maintained for at least three passages following recovery from frozen stocks before use in toxicity experiments. For viability measurements, 15,000 cells per well were seeded into clear 96-well plates in 100 μl growth medium and allowed to adhere for 24 h. Cells were subsequently treated with insulin aggregation samples and incubated for an additional 48 h. Treatment conditions included FAIM-S alone (5 or 20 μM), insulin alone (80 μM), and insulin aggregation samples diluted to final insulin concentrations of 5, 10, 20, or 40 μM in the presence of either 5 or 20 μM FAIM-S. FAIM-S and insulin treatment solutions were prepared fresh before each experiment. Cell viability was assessed using the CyQUANT XTT Cell Viability Assay (Thermo Fisher Scientific, X12223) according to the manufacturer’s instructions. XTT reagent was added to each well, and plates were incubated for 4 h at 37 °C before absorbance measurements were acquired at 450 nm with a reference wavelength of 650 nm. Background-corrected absorbance values (A − A) were calculated following subtraction of cell-free blank measurements and normalized to the buffer-only control. Data are presented from three independent replicates. Statistical significance was assessed by one-way analysis of variance (ANOVA) followed by Tukey’s honestly significant difference (HSD) post hoc test using MATLAB (MathWorks). Pairwise comparisons were performed relative to the insulin-only treatment group. Differences were considered significant at *p* < 0.05.

### NMR spectroscopy

Samples contained 80 μM insulin, 10 mM sodium phosphate, 150 mM NaCl, and 0, 5, 10, or 20 μM FAIM-S at pH 3.0. A FAIM-S-only control (10 μM) was prepared under identical buffer conditions. Aliquots were collected from aggregation reactions at the indicated time points, flash-frozen, and stored at −80 °C until analysis. Before data acquisition, samples were thawed and supplemented with 10% D□O for field locking. NMR experiments were performed on a 700 MHz Bruker spectrometer equipped with a 5 mm TCI cryoprobe. Samples were maintained on ice before loading into the probe. One-dimensional ^1^H NMR spectra were acquired with 1024 scans, a recycle delay of 1.25 s, and WATERGATE water suppression. Spectra were collected at 37 °C, and samples remained in the magnet at this temperature between successive acquisitions.

### Structural prediction with AlphaFold3

Models of FAIM with insulin A and B chains were constructed using the AlphaFold3 server (https://alphafoldserver.com/welcome). The resulting model left insulin largely intact, despite lacking explicit information about disulfide bonds or the folded insulin structure and was used as the starting point for development of a pH 3.0 structural model of FAIM:Insulin (A+B). From the resulting mmCIF files, the FAIM structure was processed with crimm (https://github.com/BrooksResearchGroup-UM/crimm/) and pyCHARMM^68^ to be consistent with a pH 3.0 environment. In this model all acidic residues (Asp and Glu) were protonated. The processed FAIM structure served as the starting point for subsequent molecular dynamics simulations. Since the disulfide bonds within and between the insulin A and B chains were not enforced in the AlphaFold3 structure prediction, although the corresponding cysteine residues were very close to their SG-SG bonded positions in the predicted structure, the crystallographic model for human insulin (pdb entry 2m1e) was processed with crimm and pyCHARMM to obtain a pH 3.0 model of insulin A and B chains with intact disulfide bonds. This model was then subjected to 320 ns of molecular dynamics with the CHARMM implicit solvent model. The results were clustered using K-means, and the structure closest to the most populated cluster centroid was chosen for subsequent modeling. This refined insulin structure was overlaid onto the AlphaFold3 FAIM–insulin complex to replace the low pH insulin model, and the resulting FAIM–insulin complex was then subjected to an additional 320 ns of pH 3.0 GBMV molecular dynamics. The most populated cluster (structure closest to the centroid) from this simulation was used in the analysis described above. All structural analyses reported here were based on this FAIM–insulin complex model at pH 3.0. The insulin-only pH 3.0 dynamics model was generated to provide a refined insulin structure under low pH conditions but was not used for downstream analyses.

## Supplementary Information

**Fig. S1.**
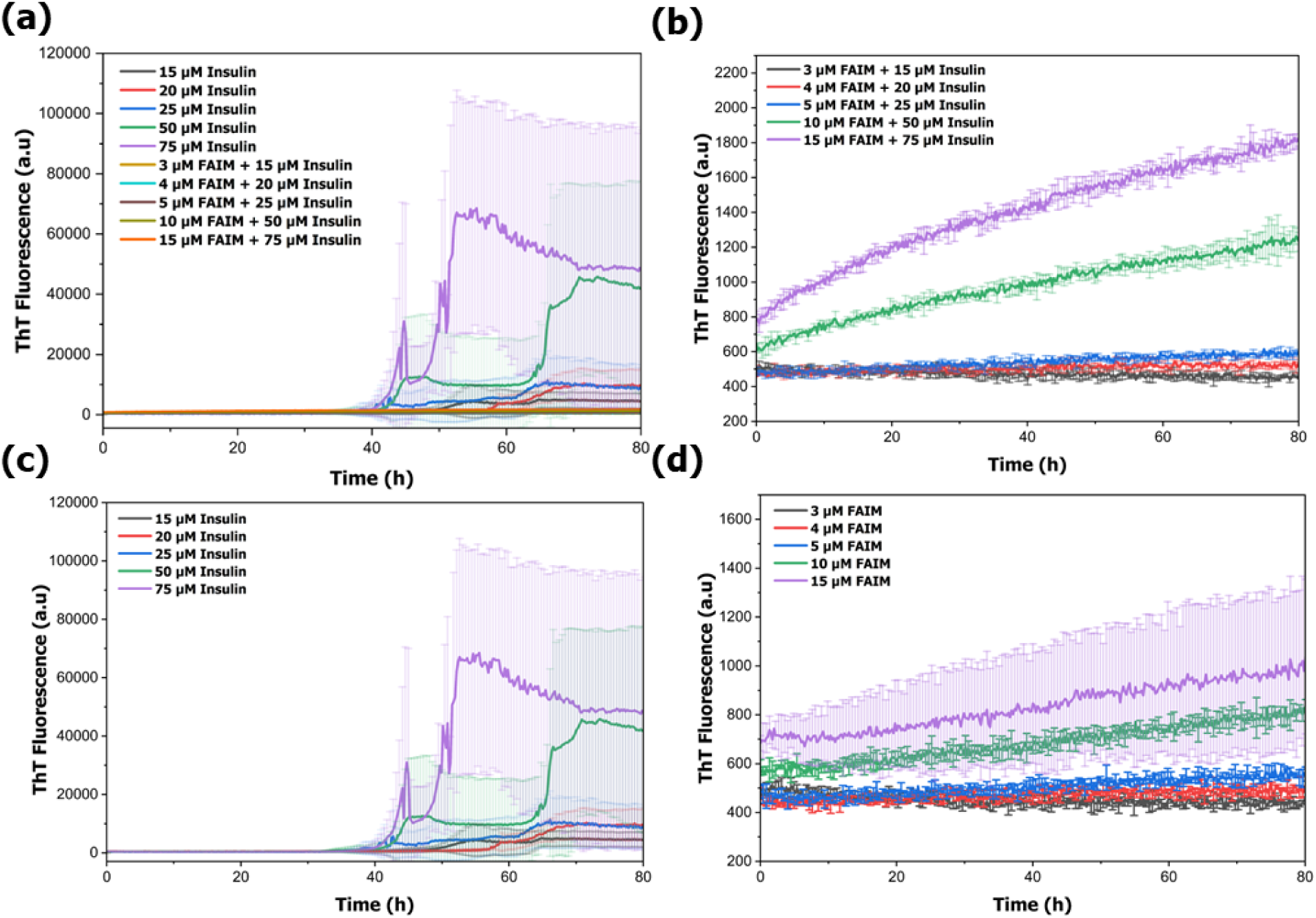
**(a)** ThT fluorescence kinetics of insulin incubated at concentrations ranging from 15 to 75 μM in the absence or presence of FAIM at a fixed FAIM:insulin molar ratio of 1:5. **(b)** Expanded view of the FAIM-containing samples shown in **(a)**. **(c)** Insulin-only samples. **(d)** FAIM-only samples. Conditions: 10 mM sodium phosphate, 150 mM NaCl, 10 μM ThT, pH 3.0, 37 °C, 700 rpm. Data represent mean ± SD from three independent experiments.

**Table S1.** BeStSel structural analysis of CD spectra of insulin co-incubated with FAIM. CD spectra were analyzed using the BeStSel online server to quantify secondary structure content of 80 μM insulin incubated with increasing concentrations of FAIM (5, 10, or 20 μM), as well as control conditions with 10 μM FAIM alone and 80 μM insulin alone. The percentages of regular and distorted helices, β-strand components (Anti1, Anti2, Anti3), turn structures, and unordered or other structures are shown.

| **Secondary Structure (%)** | **5 μM FAIM + 80 μM Insulin** | **10 μM FAIM + 80 μM Insulin** | **20 μM FAIM + 80 μM Insulin** | **10 μM FAIM** | **80 μM Insulin** |
| --- | --- | --- | --- | --- | --- |
| Helix 1 (regular) | 6.0 | 1.0 | 3.0 | 0.0 | 0.0 |
| Helix 2 (distorted) | 5.0 | 1.0 | 3.0 | 0.0 | 0.0 |
| Anti1 (left-twisted) | 1.0 | 2.0 | 2.0 | 1.0 | 1.0 |
| Anti2 (relaxed) | 20.0 | 15.0 | 14.0 | 17.0 | 15.0 |
| Anti3 (right-twisted) | 12.0 | 17.0 | 16.0 | 19.0 | 19.0 |
| Parallel | 0.0 | 0.0 | 0.0 | 0.0 | 0.0 |
| Turn | 14.0 | 16.0 | 16.0 | 17.0 | 16.0 |
| Others | 47.0 | 49.0 | 48.0 | 47.0 | 48.0 |

**Fig. S2.**
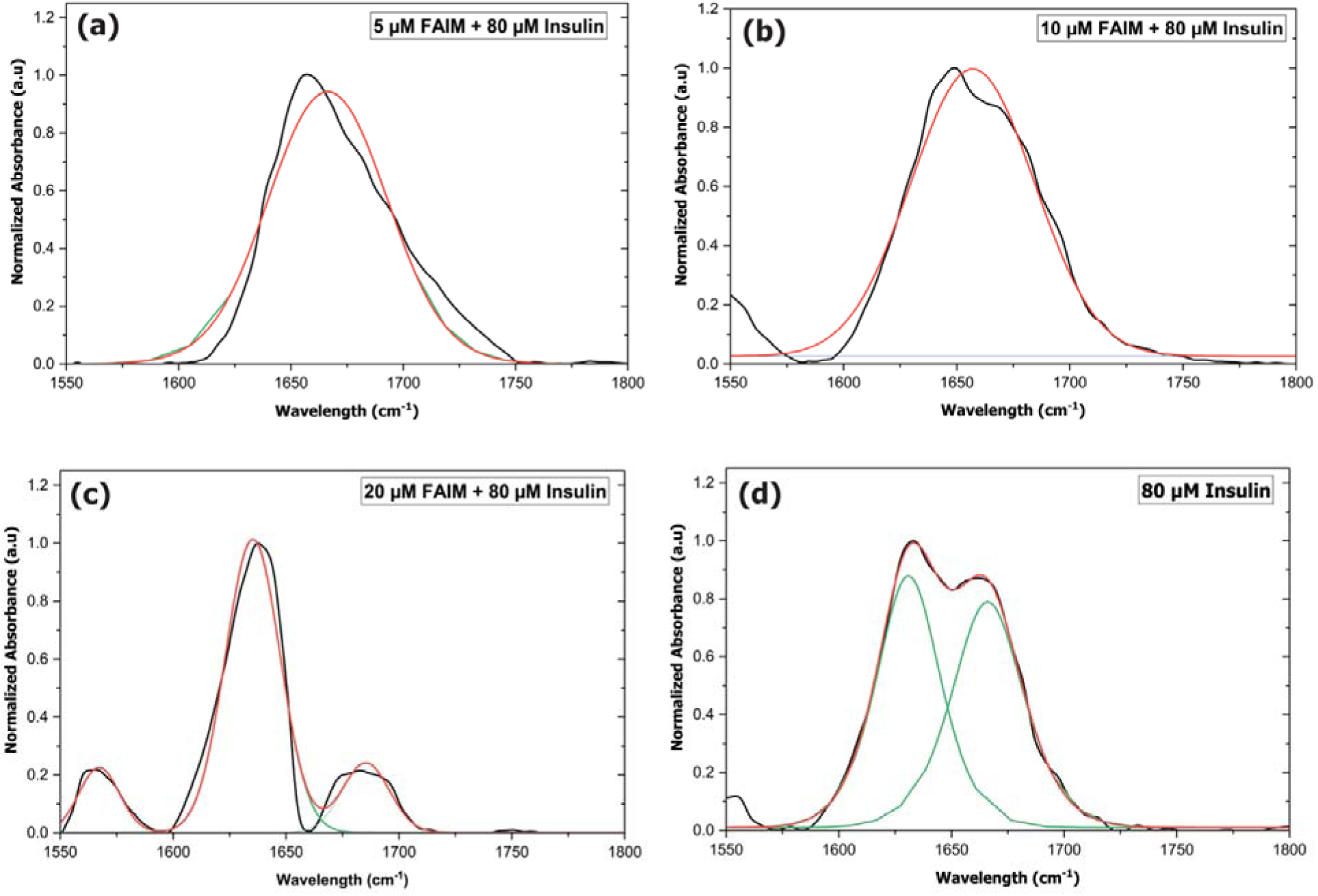
Gaussian deconvolution of FTIR spectra for insulin aggregates co-incubated with FAIM. Samples contained 80 µM insulin incubated alone or with 5, 10, or 20 µM FAIM, and were aggregated for 48 h under shaking at 37°C. FTIR spectra were collected in the amide I region (1550–1800 cm□^1^), where absorbance corresponds to protein secondary structure. Peak fitting was performed in OriginPRO using a Gaussian model to resolve overlapping spectral features. Raw spectra (black), global Gaussian fits (red), and individual secondary structure components (green) are shown. FAIM co-incubation induced shifts in the β-sheet–associated peak (∼1625–1640 cm□^1^) and a redistribution of intensity, indicating reduced β-sheet content and altered fibril morphology.

**Table S2.**
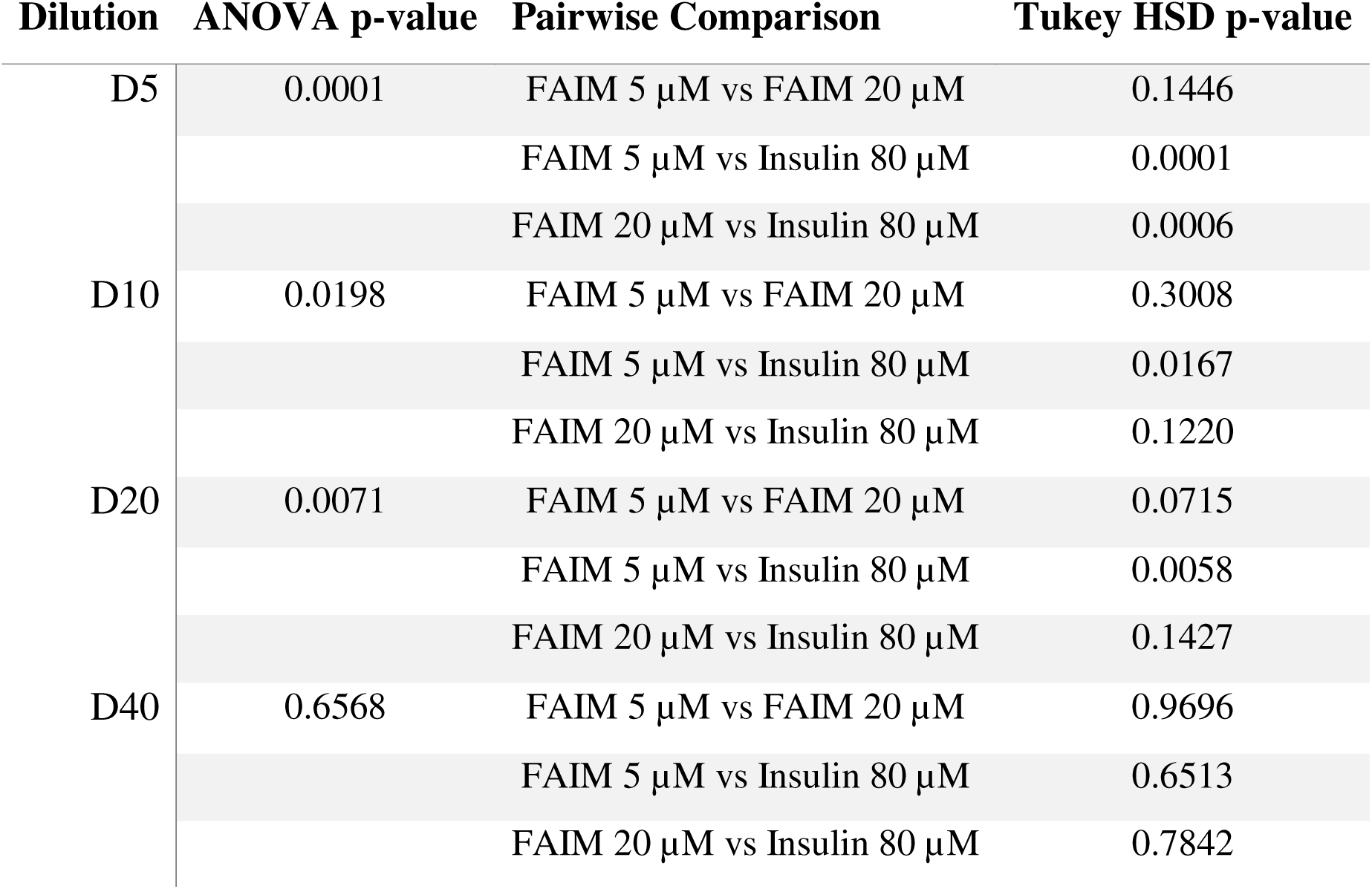
One-way ANOVA and Tukey HSD pairwise comparisons for each dilution condition (D5–D40). Values indicate p-values for overall group differences and post-hoc pairwise tests between FAIM-treated and insulin-only samples.

| Dilution | ANOVA p-value | Pairwise Comparison | Tukey HSD p-value |
| --- | --- | --- | --- |
| D5 | 0.0001 | FAIM 5 μM vs FAIM 20 μM | 0.1446 |
| | | FAIM 5 μM vs Insulin 80 μM | 0.0001 |
| | | FAIM 20 μM vs Insulin 80 μM | 0.0006 |
| D10 | 0.0198 | FAIM 5 μM vs FAIM 20 μM | 0.3008 |
| | | FAIM 5 μM vs Insulin 80 μM | 0.0167 |
| | | FAIM 20 μM vs Insulin 80 μM | 0.1220 |
| D20 | 0.0071 | FAIM 5 μM vs FAIM 20 μM | 0.0715 |
| | | FAIM 5 μM vs Insulin 80 μM | 0.0058 |
| | | FAIM 20 μM vs Insulin 80 μM | 0.1427 |
| D40 | 0.6568 | FAIM 5 μM vs FAIM 20 μM | 0.9696 |
| | | FAIM 5 μM vs Insulin 80 μM | 0.6513 |
| | | FAIM 20 μM vs Insulin 80 μM | 0.7842 |

**Fig. S3.**
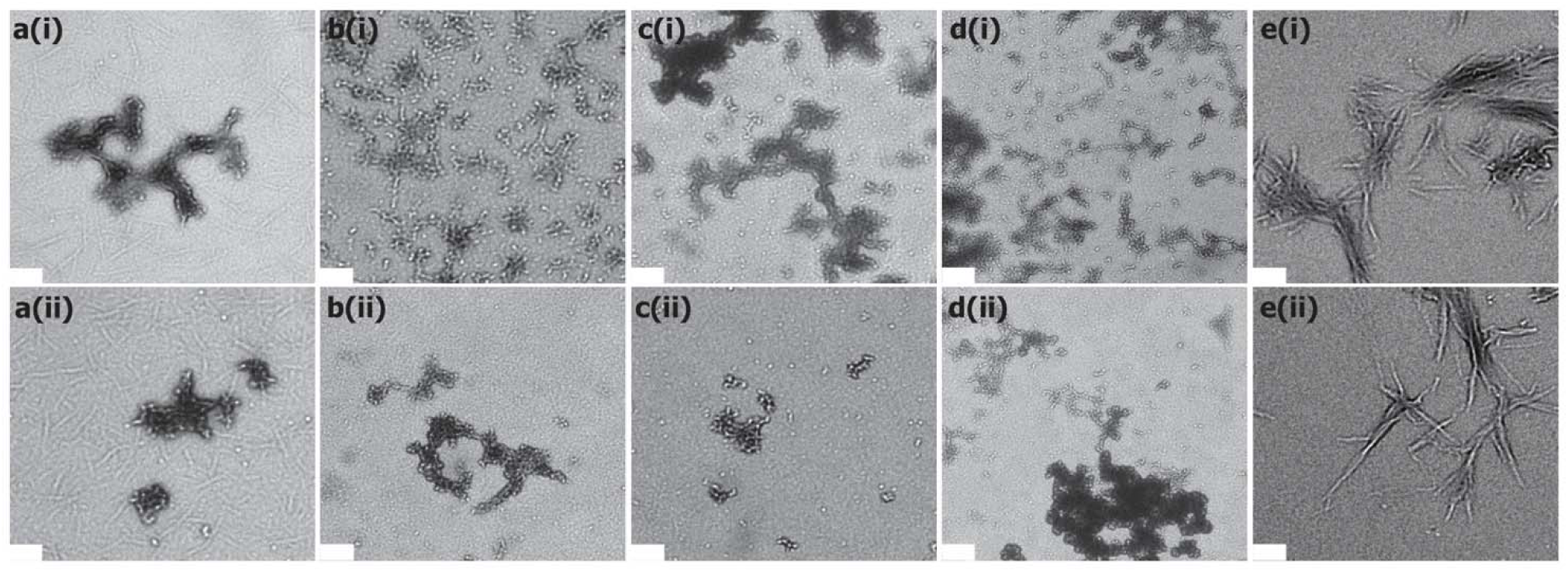
TEM images of FAIM and insulin aggregates. **(a-c)** Insulin co-incubated with FAIM at concentrations of 5 µM, 10 µM, and 20 µM, respectively. **(d)** FAIM alone. **(e)** Insulin alone. For each condition, panels (i) and (ii) represent images captured from different regions of the same grid to illustrate variability across the sample. Samples were taken after 48 h of incubation. The scale bar (in white) represents 200 nm.

**File S1.** MATLAB code used to perform ANOVA and Tukey HSD comparisons for the FAIM + insulin dilution series (FAIM_Insulin_Cytotoxicity_Analysis.m).

## Acknowledgements

The authors thank Dr. Hiroaki Kaku for providing the plasmid used for generating FAIM protein. They also acknowledge Ms. Isabella Brown for her contribution to the graphical abstract. This study was supported by NIDDK, NIH (DK132214).

## Author Contributions

Dana Wolfe: Writing – original draft (lead); Writing – review and editing (lead); Investigation (lead); Formal Analysis (lead); Methodology (supporting); Validation (lead); Visualization (lead);

Jhinuk Saha: Writing – review and editing (supporting); Investigation (supporting); Formal Analysis (supporting); Methodology (lead); Validation (supporting)

Joshua Mitchell: Investigation (supporting); review and editing (supporting)

Sam McCalpin: Investigation (supporting); Formal Analysis (supporting); Visualization (supporting)

Michael Gutknecht: Investigation (supporting)

Charles L. Brooks III: Software (lead); Methodology (supporting)

Thomas L. Rothstein: Conceptualization (supporting); review and editing (supporting); Funding Acquisition (supporting); Resources (supporting)

Ayyalusamy Ramamoorthy: Conceptualization (lead); Funding Acquisition (lead); Methodology (lead); Supervision (lead); Writing – review and editing (supporting); Resources (lead)

